# Taxonomy of interventions at academic institutions to improve research quality

**DOI:** 10.1101/2022.12.08.519666

**Authors:** Alexandra R Davidson, Ginny Barbour, Shinichi Nakagawa, Alex O. Holcombe, Fiona Fidler, Paul P Glasziou

## Abstract

Research institutions and researchers have become increasingly concerned about poor research reproducibility and replicability, and research waste more broadly. Research institutions play an important role and understanding their intervention options is important. This review aims to identify and classify possible interventions to improve research quality, reduce waste, and improve reproducibility and replicability within research-performing institutions.

Taxonomy development steps: 1) use of an exemplar paper of journal-level research quality improvement interventions, 2) 2-stage search in PubMed using seed and exemplar articles, and forward and backward citation searching to identify articles evaluating or describing research quality improvement, 3) elicited draft taxonomy feedback from researchers at an open-sciences conference workshop, and 4) cycles of revisions from the research team.

The search identified 11 peer-reviewed articles on relevant interventions. Overall, 93 interventions were identified from peer-review literature and researcher reporting. Interventions covered before, during, and after study conduct research stages and whole of institution. Types of intervention included: Tools, Education & Training, Incentives, Modelling & Mentoring, Review & Feedback, Expert involvement, and Policies & Procedures. Identified areas for research institutions to focus on to improve research quality and for further research includes improving incentives to implement quality research practices, evaluating current interventions, encourage no- or low-cost/high-benefit interventions, examine institution research culture, and encourage mentor-mentee relationships.

## Introduction

Over the past decade, the problems of research waste and the reproducibility crisis have been extensively documented.(1-4) A 2014 series in the Lancet demonstrated that approximately 85% of biomedical research goes to waste through the combination of poor design, non-publication, and poor reporting, (1) with a similar percent recently reported for ecology research.(5) Studies in disciplines as diverse as economics, cancer biology, psychology, machine learning, ecology, and social sciences have found disappointingly low reproducibility and replicability.(1-4)

Low reproducibility means that original protocol, materials or data sets may not be available to conduct analysis to reproduce the results, and low replicability relates to the inability to re-conduct or re-conduct well an entire study or experiment, regardless of whether the results replicate.(6) The two approaches, reproducibility and replicability, exist on a spectrum from ‘direct’ following the original methods strictly, to ‘conceptual’ where researchers may selectively alter aspects of the original methods to test for robustness and generalisability.(7, 8) Both are important to reducing research waste and improving overall research practice quality.

Poor research reproducibility and replicability is partly attributable to flaws in study design and partly to incomplete or poor documentation of research processes. The flow-on effects impact research end-users such as industries that utilize research to facilitate practice. Many of these problems are avoidable and might be reduced with sustained interventions at the research systems level. The key stakeholders in improving the research system to improve quality are the research funders and research institutions.

What might research institutions do to improve the quality and reproducibility of their research? This work builds on a previous taxonomy of interventions for journals and publishers developed by Blanco et al in their scoping review of interventions to improve adherence to reporting guidelines in health research, which classifies interventions by the type of intervention and by the research stage.(9) The current taxonomy expands the behaviour change categories used by Blanco, drawing on Michie’s behaviour change wheel which covers Training, Incentivisation, Modelling, Persuasion, Education and Coercion.(10) Using this approach is useful, not just for identifying existing interventions, but also to identify where there are gaps. We are not aware of any previous study that classifies interventions to improve quality and increase reproducibility at institutions.

This review aims to identify and classify possible interventions to improve research quality, reduce waste, and improve reproducibility and replicability within research-performing institutions. We also identified studies that assessed the interventions.

## Methods

Research institutions were inclusive of academic institutions such as universities, government research institutes, and privately funded institutions. Within institutions, interventions could occur at a range of levels, from individual actions to department and whole of institution levels, including policy changes.

The interventions could be training or education, institutional incentives or regulations, or provision of infrastructure and tools; the only requirement was that the intervention must be aimed at some aspect of reducing research waste, improving quality, or improving reproducibility. For example, interventions aimed at better study design or better conduct of research, increased or timelier publication of research, better reporting of research, including better “open science” such as the provision of protocols and other research process details, and research data would all be includable.

### The Search

Because the potential range of potential interventions and terms used was broad and unknown, we used a 2-stage process for the search. Stage 1 used a set of seed articles and reviews identified by the authors from a preliminary search which identified several articles including a review of journal interventions.(9) We then used a forward and backward citations search of this set of articles to widen the pool of potential articles. Stage 2 then conducted a word frequency analysis on these eligible articles to identify key terms to build a search strategy for the full database searches.

This Stage 1 search identified a key review article on interventions to improve adherence to reporting guidelines for journals, which included a suggested taxonomy.(9) We then drafted a potential taxonomy and used the other interventions identified from the searches to test and modify this proposed intervention taxonomy.

Next, we sought input from others for further examples and on the taxonomy. To ensure the taxonomy was reflective of research practice in institutions, we invited possible end-users to assist in co-design. During the 2021 AIMOS Association for Interdisciplinary Meta-Research and Open Science Conference (https://www.ivvy.com.au/event/aimos2021), we held a workshop with approximately 40 participants to further refine the draft taxonomy. Workshop participants included researchers at different career levels, ranging from PhD students to professors.

Briefly, the steps of the workshop process were

1. List any interventions you have conducted, attended, or heard of.
2. Map these interventions onto the taxonomy using a Google Doc accessible to all participants (Note: if they do not fit, then put them into the second list)
3. Discussion of interventions that do not fit the proposed taxonomy (do these warrant a change to the taxonomy?)
4. General Discussion on next steps

Following the workshop, we used the participant input to develop the revised taxonomy, collect further potential examples and revised the taxonomy again.

## Results

### The taxonomy

Interventions were first classified according to the research stage of their implementation: before study conduct, during study conduct, and after study conduct. Research stages were then further subclassified into education; grant writing; protocol writing; research conduct & analysis; manuscript writing; manuscript submission; and post-publication. Table 1 highlights which type of behaviour change interventions, as classified by Michie’s behaviour wheel, are represented at each research stage.

**Table 1:**
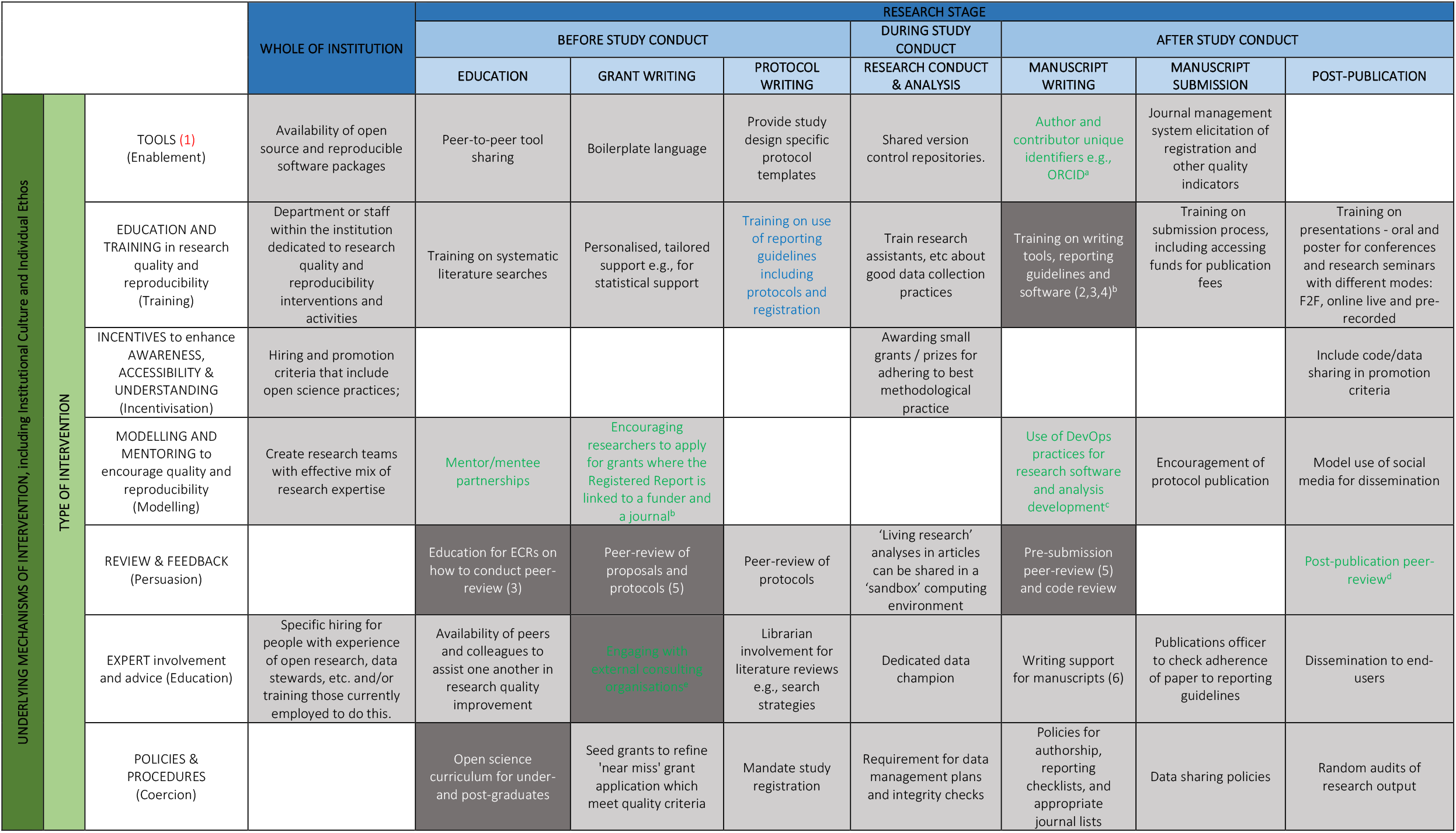
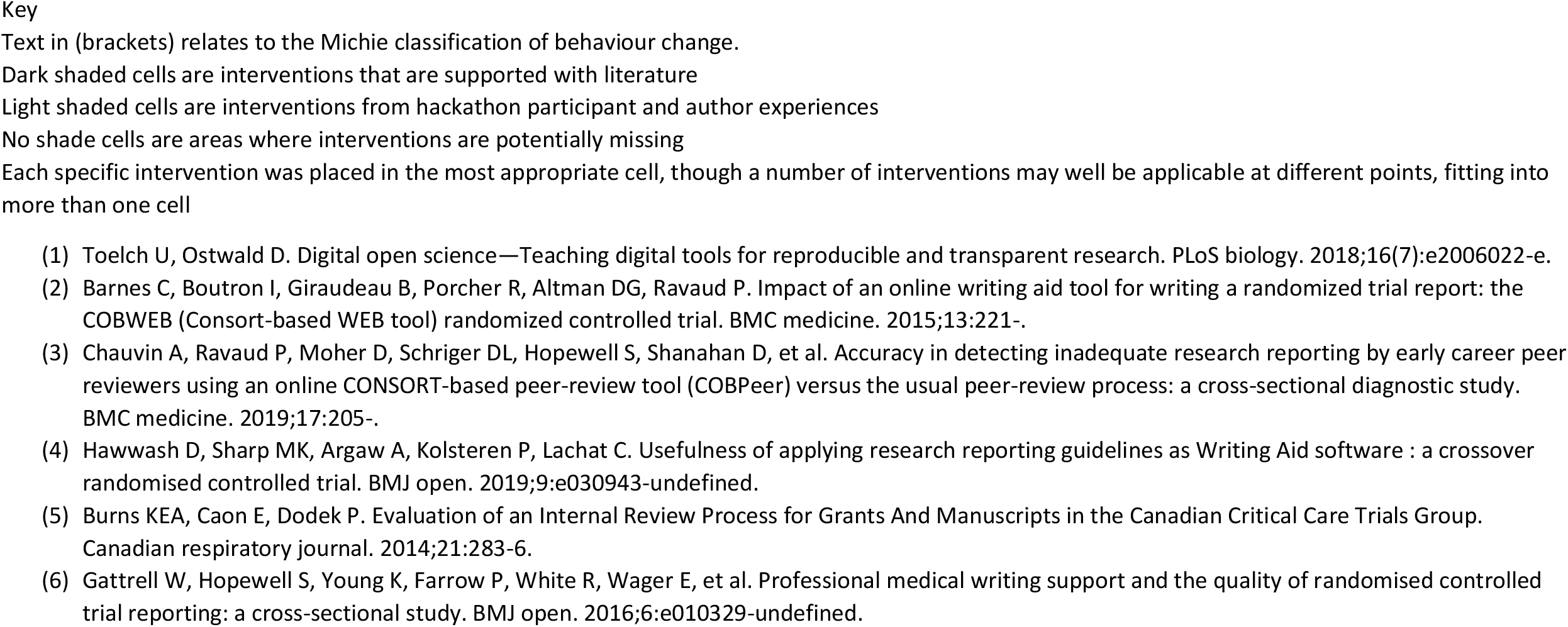
Outline of classification of interventions and their relationship to research stage – condensed version of taxonomy

|  |  |  | WHOLE OF INSTITUTION | RESEARCH STAGE |  |  |  |  |  |  |
| --- | --- | --- | --- | --- | --- | --- | --- | --- | --- | --- |
|  |  |  |  | BEFORE STUDY CONDUCT |  |  | DURING STUDY CONDUCT | AFTER STUDY CONDUCT |  |  |
|  |  |  |  | EDUCATION | GRANT WRITING | PROTOCOL WRITING | RESEARCH CONDUCT & ANALYSIS | MANUSCRIPT WRITING | MANUSCRIPT SUBMISSION | POST-PUBLICATION |
| UNDERLYING MECHANISMS OF INTERVENTION, including Institutional Culture and Individual Ethos | TYPE OF INTERVENTION | TOOLS (1)<br>(Enablement) | Availability of open source and reproducible software packages | Peer-to-peer tool sharing | Boilerplate language | Provide study design specific protocol templates | Shared version control repositories. | Author and contributor unique identifiers e.g., ORCID<sup>a</sup> | Journal management system elicitation of registration and other quality indicators |  |
|  |  | EDUCATION AND TRAINING in research quality and reproducibility (Training) | Department or staff within the institution dedicated to research quality and reproducibility interventions and activities | Training on systematic literature searches | Personalised, tailored support e.g., for statistical support | Training on use of reporting guidelines including protocols and registration | Train research assistants, etc about good data collection practices | Training on writing tools, reporting guidelines and software (2,3,4)<sup>b</sup> | Training on submission process, including accessing funds for publication fees | Training on presentations - oral and poster for conferences and research seminars with different modes: F2F, online live and pre-recorded |
|  |  | INCENTIVES to enhance AWARENESS, ACCESSIBILITY & UNDERSTANDING (Incentivisation) | Hiring and promotion criteria that include open science practices; |  |  |  | Awarding small grants / prizes for adhering to best methodological practice |  |  | Include code/data sharing in promotion criteria |
|  |  | MODELLING AND MENTORING to encourage quality and reproducibility (Modelling) | Create research teams with effective mix of research expertise | Mentor/mentee partnerships | Encouraging researchers to apply for grants where the Registered Report is linked to a funder and a journal<sup>b</sup> |  |  | Use of DevOps practices for research software and analysis development<sup>c</sup> | Encouragement of protocol publication | Model use of social media for dissemination |
|  |  | REVIEW & FEEDBACK (Persuasion) |  | Education for ECRs on how to conduct peer-review (3) | Peer-review of proposals and protocols (5) | Peer-review of protocols | 'Living research' analyses in articles can be shared in a 'sandbox' computing environment | Pre-submission peer-review (5) and code review |  | Post-publication peer-review<sup>d</sup> |
|  |  | EXPERT involvement and advice (Education) | Specific hiring for people with experience of open research, data stewards, etc. and/or training those currently employed to do this. | Availability of peers and colleagues to assist one another in research quality improvement | Engaging with external consulting organisations<sup>e</sup> | Librarian involvement for literature reviews e.g., search strategies | Dedicated data champion | Writing support for manuscripts (6) | Publications officer to check adherence of paper to reporting guidelines | Dissemination to end-users |
|  |  | POLICIES & PROCEDURES (Coercion) |  | Open science curriculum for under- and post-graduates | Seed grants to refine 'near miss' grant application which meet quality criteria | Mandate study registration | Requirement for data management plans and integrity checks | Policies for authorship, reporting checklists, and appropriate journal lists | Data sharing policies | Random audits of research output |

“Whole of institution” was included as an additional category, separate to the research stages, as some interventions relate to two or more stages of research or support overall research practices in that institution. Similarly, “Institutional Culture and Individual Ethos” was added to the taxonomy to highlight the influence of the culture of the institution including their overall research aims and mission, and those that work in the institution and their individual ethos, values, and attitudes towards research practices.

Table 1 gives examples of the interventions identified. The full set of interventions are displayed in Supplementary File 1. Overall, we identified 93 different possible interventions.

The types of intervention varied widely, from whole of institution policies – such as modifying hiring and promotion criteria to emphasise rigorous research design, reproducibility, and transparency, to highly specific departmental-level interventions such as developing mentor-mentee relationships. Most interventions are applicable to researchers at all levels of experience. Several interventions are specific to particular areas of research e.g., registration of clinical trials in healthcare, but others, such as mentoring, or journal clubs are relevant to multiple disciplines. We did not subclassify by disciplines.

In reviewing the taxonomy, several themes emerged. Many of the interventions require a substantial and long-term investment in people– e.g., hiring of specific experts; training of research assistants and others on data collection methods and techniques; co-design with patients and public/end-users. Though ad hoc seminars have value, most of the interventions require individuals or teams or to be embedded in institutions, even if the intervention is to provide “just in time” advice. For example, a publications officer to check adherence of papers to reporting guidelines would have to be well established for them to be able to provide on the spot advice at a time of need.

Education of researchers and research support staff can happen by a variety of formal and informal methods. There was some suggestion that some of the training had to be compulsory e.g., included in the curriculum for undergraduate and postgraduate research training. However, there was also specific recognition of the role of informal networks including peer-to-peer learning and mentor-mentee relationships. We note that mentor-mentee learnings can be in both directions, as more junior staff are sometimes the instigators of novel research practices learned during their research skill development. As training in undergraduate and postgraduate programs are constantly changing, more experienced researchers can be exposed to this by mentoring a student or Early Career Researcher.

There were surprisingly few technical interventions suggested. Most of these also included an element of human intervention e.g., use of pull-requests and code commentary by collaborators and/or external peers on shared codebases. Notably one of the technical interventions was to cease subsidising (through purchase of site licenses) the use of certain software programs (e.g., statistical and spreadsheet) that are not conducive to reproducibility, and to instead promote and encourage the use of open science source software and practices.

Surprisingly, the “incentives” row has more empty cells than the other rows. This indicates either a lack of awareness of participants involved in reviewing the taxonomy, and/or a lack of incentives being implemented and available at research institutions to encourage researchers to participant in quality research practices.

Following the classification, we searched for papers that had described and/or assessed these interventions. During the search processes, eleven articles evaluating interventions were found. All the interventions that had been assessed were in the manuscript and grant writing, or education phase of research (Table 2).

**Table 2:** All primary literature found on interventions that aim to improve research quality and reproducibility at the institutional level.

| **Citation No.** | **Author, year** | **Population** | **Aim** | **Type of Intervention** | **Research Phase** |
| --- | --- | --- | --- | --- | --- |
| **(11)** | Barnes, 2015 | Masters and doctoral students in public health and medical research | To evaluate the impact of an online writing aid tool on the completeness of reporting of two-arm parallel-group RCTs evaluating pharmacologic and non-pharmacologic interventions | Training | Manuscript writing |
| **(12)** | Boschen, 2021 | APA full-text journals | Evaluation of JATSdecoder as an automated tool to facilitate checking of reported statistical results for consistency and completeness | Tools | Manuscript writing |
| **(13)** | Burns, 2014 | Authors and reviewers of the Canadian Critical Care Trial group for grants and manuscripts | To formally evaluate authors' and reviewers' perceptions of internal peer review before journal submission | Review & Feedback | Manuscript Writing |
| **(14)** | Chauvin, 2019 | ECRs (although at the journal-level, is translatable?) | To evaluate the accuracy in identifying inadequate reporting in RCT reports by early career researchers (ECRs) using an online CONSORT-based peer-review tool (COBPeer) versus the usual peer-review process. | Training Review & Feedback | Manuscript writing Education |
| **(15)** | Gattrell, 2016 | Authors of RCTs | To examine the relationship between medical writing support and quality and timeliness of reporting of randomised controlled trial results | Expert Involvement and Advice | Manuscript writing |
| **(16)** | Hawwash, 2019 | Doctoral and postdoctoral researchers | To assess the intention to use a Writing Aid software, which integrates four research reporting guidelines (Consolidated Standards of Reporting Trials, Preferred Reporting Items for Systematic Reviews and Meta- Analyses, Strengthening the Reporting of Observational Studies in Epidemiology and Strengthening the Reporting of Observational Studies in Epidemiology- nutritional epidemiology) and their Elaboration & Explanation (E&E) documents during the write- up of research in Microsoft Word compared with current practices | Training | Manuscript writing |
| **(17)** | Hirschey, 2019 | Advance Practice Nurses doing a Doctor of | To enhance APNs' writing skills with a series of online modules, a workshop, and manuscript checklist. | Training | Manuscript writing |
|  |  | Nursing Practice program. |  |  |  |
| (18) | Nuijten, 2020 | Full text of APA journals | To describe the *statcheck* tool and provide an example of use in a meta-analysis | Tools | Manuscript writing |
| (19) | Shanahan, 2017 | Authors of speciality medical research journals | To investigate whether a decision tree tool made available during the submission process facilitates author identification of the relevant reporting guideline. | Policy & Procedures | Manuscript Writing |
| (20) | Struthers, 2021 | Authors submitting to BMJ Open | To provide an outline of the reporting guideline identification tool, GoodReports.org, and to describe user experience and behaviour of using the tool inside and outside of manuscript submission to a journal. | Tools | Manuscript Writing |
| (21) | Toelch, 2018 | University-level research course students | To evaluate an introductory digital tools course that guides students towards a reproducible science workflow – including research transparency and reproducibility. | Policy & Procedures | Education |

## Discussion

To improve institutional interventions that might improve research quality, an understanding of the range and types of interventions is vital. Based on work from Blanco et al (2019) in their scoping review of interventions to improve adherence to reporting guidelines in health research we have developed a taxonomy of possible interventions to improve research quality and reproducibility within institutions. At the institutional level, interventions are possible at all stages of research and for each stage of research there are several possible mechanisms of intervention.

Through an iterative crowdsourced process, we identified interventions that the authors or the hackathon participants had experienced, that they had seen conducted in their or others’ institutions, or which they wanted to see. Very few of the interventions have been evaluated. In several areas where interventions are possible none were identified, or the interventions suggested are only aspirational at this point.

Research quality has become a much-discussed topic in Australia, and internationally, but there is no systematic approach to improving research quality, especially regarding what interventions are needed at institutions. In Australia, research quality is most prominently assessed through the Excellence in Research for Australia (ERA) process. However, ERA assesses research outputs, not any part of the research process. An increased national focus on research quality more widely through the work of the NHMRC’s Research Quality Steering Committee (RQSC) established in 2018. In 2019, the RQSC oversaw a survey of Australian Research institutions and researchers.(22) Key opportunities identified from that survey relevant to interventions at institutions were the need for effective training and mentorship (especially of junior researchers) about responsible research practice; addressing factors that adversely affect research quality, such as poor research practices; promoting positive initiatives and processes rather than competition where possible; and encouraging more rigorous reproducibility procedures.

A recent Australian Chief Scientist declared a need to “shift from quantity to quality” and to challenge the status quo of “a passive apprenticeship system” of researcher training.(23, 24)

In the recently released National Research Infrastructure RoadMap, the importance of research quality is recognised in specific, limited areas, primarily data e.g., “An important driver for maintaining quality research output is Australia’s ability to generate and analyse data as well as improving the digital skills of researchers”.(22, 25)

On a global level, the United Nations Educational, Scientific and Cultural Organization, developed their Recommendation on Open Science in 2021.(26) Their recommendations align with the taxonomy in this paper as they recommend interventions related to institutions, including open scientific publications, research data, educational resources, source software and source code, and hardware. Open scientific publications relate to the ‘Education and Training’ – ‘Manuscript Submission’ cell – training on how to access funds for publication fees for open access journals. Open research data relates to many of the examples provided in the ‘Research Conduct & Analysis’ column of the taxonomy, including shared-version control repositories, and data dictionaries. Open educational resources, includes the examples in the ‘Education and Training’ row of the taxonomy, including the availability of training sessions to have hybrid delivery. Open-source software and source code, includes examples outlined in the ‘Tools’ row. Lastly, Open hardware relates to ‘training manuals/data collection protocol, including use of equipment’ in the ‘Research Conduct & Analysis’ – ‘Education and Training’ cell.

There have been other classifications of potential interventions, the Michie behaviour change wheel and the Nosek pyramid.(3, 10) Both of those classifications align with ours in that they range from interventions that are simply a change in the environment – our “tools”, Nosek’s “make it easy”, Michie’s “enablement” through to required actions - our “policies and procedures”, Nosek’s “make it required”, Michie’s “coercion”. What the classifications all demonstrate is the need for a range of approaches. By mapping interventions to specific research stage and interventions type, we have demonstrated the range of possible interventions, where there are gaps and especially the relative lack of assessment of these interventions.

### Limitations

The interventions in the taxonomy we present are not a comprehensive list of all possible interventions. We did not assess adherence to any of the interventions or examine their effectiveness, except where there were previously published papers. As the participants and authors were largely or exclusively from the sciences, the list of interventions may not include interventions within Humanities and Social Sciences.

A strength of this research is the use of both published peer-reviewed literature and end-user engagement to develop the taxonomy.

### Future Directions

Given most interventions outlined in the taxonomy have not been evaluated for their impact on research quality and reproducibility, there is a clear need for more institutional interventions be evaluated. Priority areas for evaluation should be those currently in common use at institutions, to assess their value. Implementation of new or different interventions could be those that are no- or low-cost, such as open access tools and software to enhance research practices, e.g., Overleaf, and JASP, and adaption of policies and the research environment to promote open science practices.

Institution culture and individual researcher ethos have a strong influence over the reproducibility, quality, and transferability of research practices. The UK Reproducibility Network encourages institutions to examine their research culture and how it may or may not be supportive of producing robust and credible research.(27) The implementation and evaluation of interventions outlined in our taxonomy should be considered along with the institution’s current culture and potential shifts that could be made to encourage and promote open science practices.

Finally, it is vital to explore the paucity of “incentive” interventions that research institutes might use. Incentivisation is important in workload models of research institutions, much like universities have incentives for their education and teaching of degrees and coursework, they need incentives for research quality. The kind of incentives depend heavily on the institutional structures and availability of resources to create or fund incentives. It is recommended that future research could be guided by this taxonomy further to identify how incentivisation of quality research practices could be better implemented.

## Acknowledgements

The authors would like to thank the AIMOS 2021 Conference Hackathon Workshop attendees who participated in providing feedback for the draft taxonomy.

---

**Supplementary File 1 – Table:**
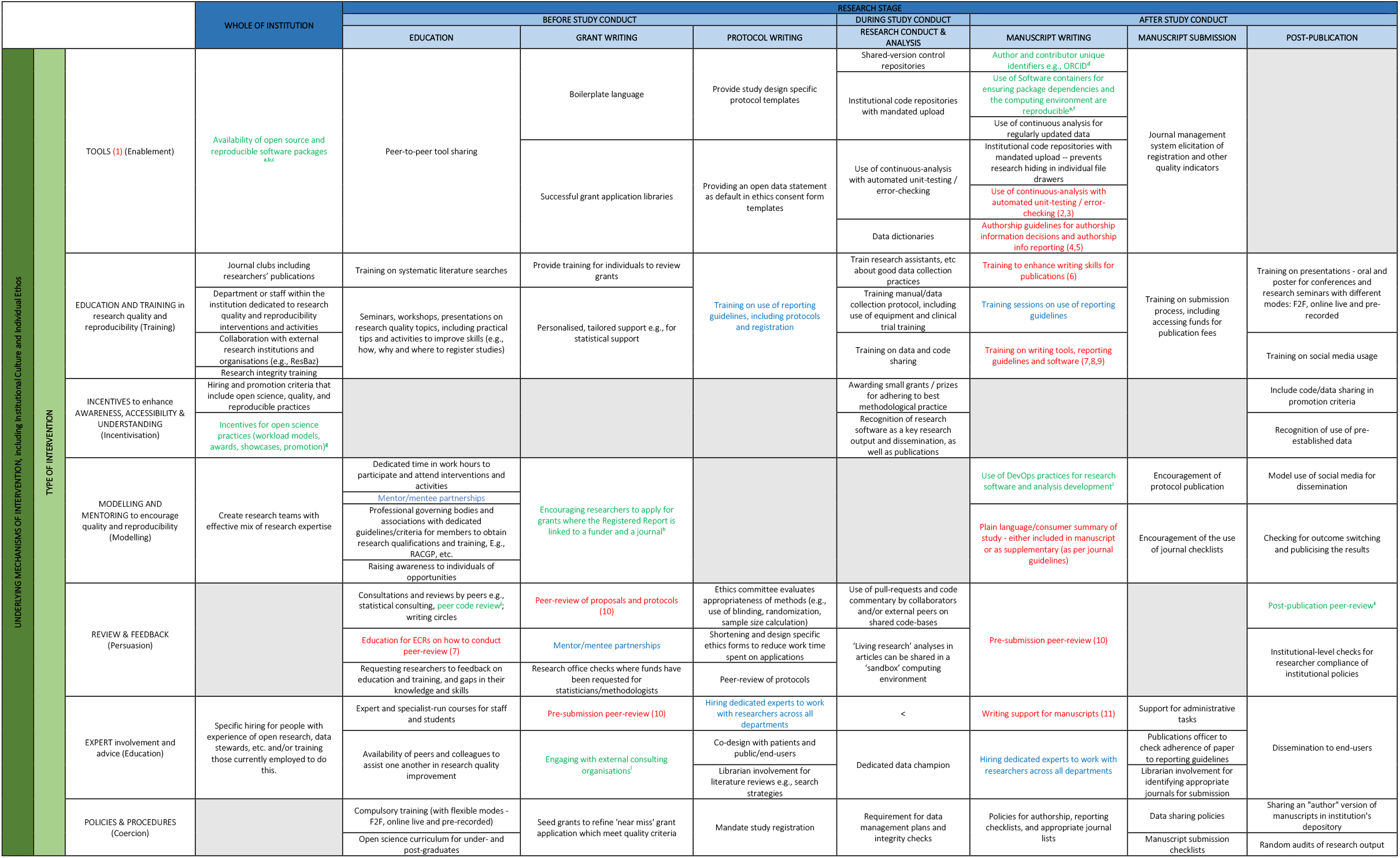
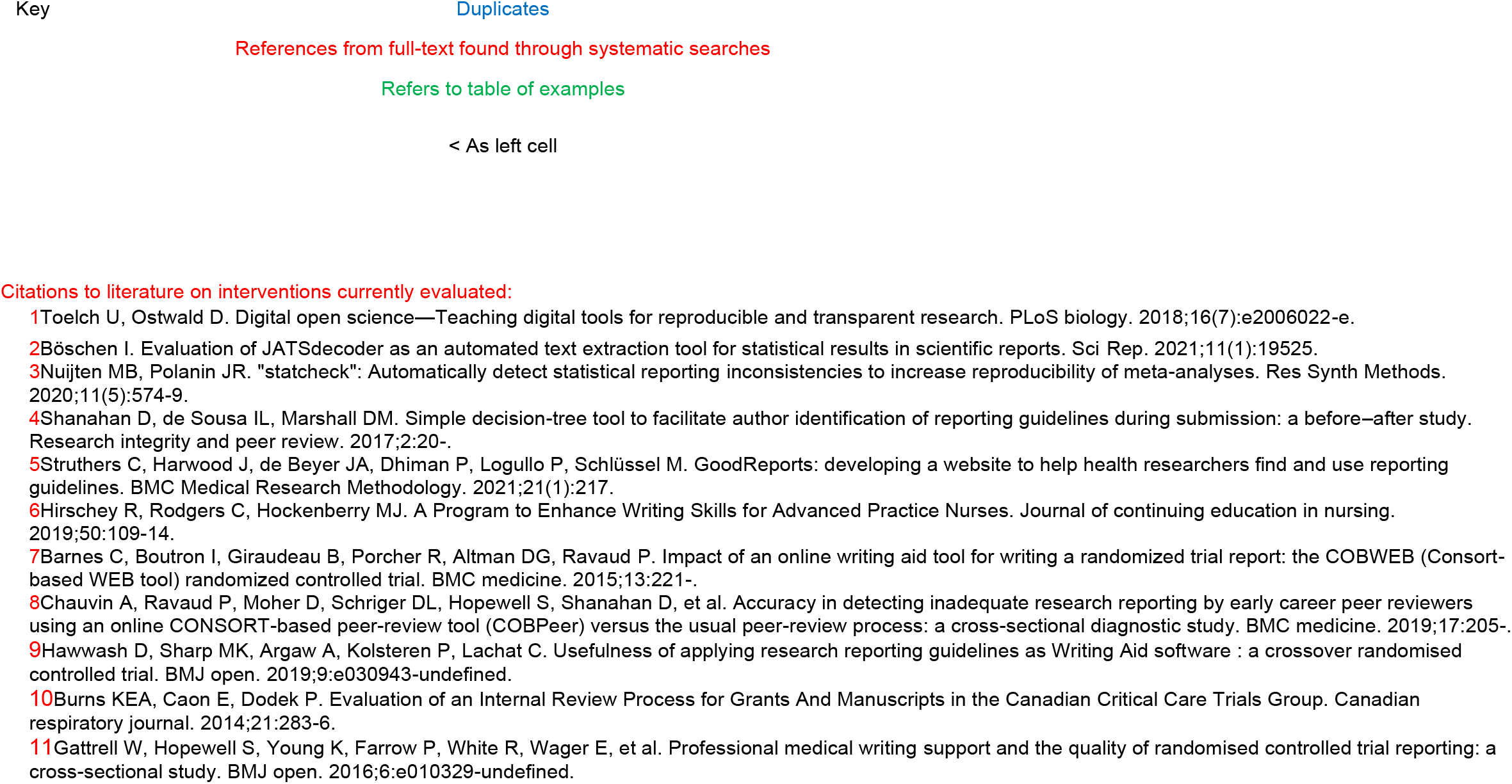
Taxonomy of interventions at academic institutions to improve research quality

|  |
| --- |
| Example Interventions |
| Tools – Whole of Institution<br>a. R – The R Project for Statistical Computing https://www.r-project.org/<br>b. Jamovi – Free and Open Statistical Software https://www.jamovi.org/<br>c. JASP – Open-source project for statistics https://jasp-stats.org/ |
| Tools - Manuscript Submission<br>d. Unique author identifier, ORCID https://orcid.org/<br>e. Overleaf https://www.overleaf.com/ Provides templates for journal articles.<br>f. Zotero https://www.zotero.org/ assists in organising, collating and sharing research. |
| Incentives – Whole of Institution<br>g. Prizes for publication of pre-registered work – OSF https://osf.io/x5w7h/wiki/06%20Leaderboard/ |
| Modelling and Mentoring and Grant Writing<br>h. “For Funders”, https://www.cos.io/initiatives/registered-reports |
| Initiatives - Manuscript Writing<br>i. Use of DevOps (https://www.atlassian.com/devops) practices for research software and analysis development:<br>- Github/GitLab for collaborative version control https://about.gitlab.com/devops-tools/github-vs-gitlab/<br>- Agile / scrum methodologies / other standard operating procedures for collaborating on code<br>- Use of Software containers for ensuring package dependencies and the computing environment are reproducible<br>- Use of continuous analysis for regularly updated data |
| Review & Feedback – Education<br>j. RAMP Rapid Review Group UK that offered rapid reviews of COVID related research which included code base review. https://epcced.github.io/ramp/previous-updates.html |
| Review & Feedback - Post-Publication<br>k. PubPeer https://pubpeer.com/static/about |
| Expert and Grant Writing<br>l. Outside Opinion https://www.outsideopinion.com.au/ |

## References

1. Glasziou P, Altman DG, Bossuyt P, Boutron I, Clarke M, Julious S, et al. Reducing waste from incomplete or unusable reports of biomedical research. Lancet. 2014;383(9913):267–76.

2. Errington TM, Iorns E, Gunn W, Tan FE, Lomax J, Nosek BA. An open investigation of the reproducibility of cancer biology research. eLife. 2014;3:e04333.

3. PSYCHOLOGY. Estimating the reproducibility of psychological science. Science. 2015;349(6251):aac4716.

4. Purgar M, Klanjscek, T., & Culina, A. Identify, quantify, act: tackling the unused potential of ecological research. 2021.

5. Purgar M, Klanjscek T, Culina A. Quantifying research waste in ecology. Nat Ecol Evol. 2022;6(9):1390–7.

6. Barba LA. Terminologies for reproducible research. arXiv preprint arXiv:180203311. 2018.

7. National Academies of Sciences E, and Medicine; Policy and Global Affairs; Committee on Science, Engineering, Medicine, and Public Policy; Board on Research Data and Information; Division on Engineering and Physical Sciences; Committee on Applied and Theoretical Statistics; Board on Mathematical Sciences and Analytics; Division on Earth and Life Studies; Nuclear and Radiation Studies Board; Division of Behavioral and Social Sciences and Education; Committee on National Statistics; Board on Behavioral, Cognitive, and Sensory Sciences; Committee on Reproducibility and Replicability in Science. Understanding Reproducibility and Replicability. Reproducibility and Replicability in Science. Washington (DC): National Academies Press (US); 2019.

8. Fidler FW, John,. Reproducibility of Scientific Results. Stanford Encyclopedia of Philosophy. 2018.

9. Blanco D, Altman D, Moher D, Boutron I, Kirkham JJ, Cobo E. Scoping review on interventions to improve adherence to reporting guidelines in health research. BMJ Open. 2019;9(5):e026589.

10. Michie S, van Stralen MM, West R. The behaviour change wheel: A new method for characterising and designing behaviour change interventions. Implementation Science. 2011;6(1):42.

11. Barnes C, Boutron I, Giraudeau B, Porcher R, Altman DG, Ravaud P. Impact of an online writing aid tool for writing a randomized trial report: the COBWEB (Consort-based WEB tool) randomized controlled trial. BMC medicine. 2015;13:221-.

12. Böschen I. Evaluation of JATSdecoder as an automated text extraction tool for statistical results in scientific reports. Sci Rep. 2021;11(1):19525.

13. Burns KEA, Caon E, Dodek P. Evaluation of an Internal Review Process for Grants And Manuscripts in the Canadian Critical Care Trials Group. Canadian respiratory journal. 2014;21:283–6.

14. Chauvin A, Ravaud P, Moher D, Schriger DL, Hopewell S, Shanahan D, et al. Accuracy in detecting inadequate research reporting by early career peer reviewers using an online CONSORT-based peer-review tool (COBPeer) versus the usual peer-review process: a cross-sectional diagnostic study. BMC medicine. 2019;17:205-.

15. Gattrell W, Hopewell S, Young K, Farrow P, White R, Wager E, et al. Professional medical writing support and the quality of randomised controlled trial reporting: a cross-sectional study. BMJ open. 2016;6:e010329–undefined.

16. Hawwash D, Sharp MK, Argaw A, Kolsteren P, Lachat C. Usefulness of applying research reporting guidelines as Writing Aid software : a crossover randomised controlled trial. BMJ open. 2019;9:e030943–undefined.

17. Hirschey R, Rodgers C, Hockenberry MJ. A Program to Enhance Writing Skills for Advanced Practice Nurses. Journal of continuing education in nursing. 2019;50:109–14.

18. Nuijten MB, Polanin JR. “statcheck”: Automatically detect statistical reporting inconsistencies to increase reproducibility of meta-analyses. Res Synth Methods. 2020;11(5):574–9.

19. Shanahan D, de Sousa IL, Marshall DM. Simple decision-tree tool to facilitate author identification of reporting guidelines during submission: a before–after study. Research integrity and peer review. 2017;2:20-.

20. Struthers C, Harwood J, de Beyer JA, Dhiman P, Logullo P, Schlüssel M. GoodReports: developing a website to help health researchers find and use reporting guidelines. BMC Medical Research Methodology. 2021;21(1):217.

21. Toelch U, Ostwald D. Digital open science—Teaching digital tools for reproducible and transparent research. PLoS biology. 2018;16(7):e2006022–e.

22. NHMRC. Research quality: National Health and Medical Research Council (NHMRC) promotes the highest quality in the research that it funds. 2020 [Available from: https://www.nhmrc.gov.au/research-policy/research-quality.

23. Finkel A. To move research from quantity to quality, go beyond good intentions. Nature. 2019;566(7744):297.

24. Finkel A. Chief Scientist calls for formal action to bake in better research practices: Australian Government 2019 [Available from: https://www.chiefscientist.gov.au/2019/02/article-chief-scientist-calls-for-formal-action-to-bake-in-better-research-practices.

25. Department of Employment and Workplace Relations. National Research Infrastructure 2022 [Available from: https://www.dese.gov.au/national-research-infrastructure.

26. United Nations Educational SaCO. UNESCO Recommendation on Open Science 2021.

27. U. K. Reproducibility Network Steering Committee. From grassroots to global: A blueprint for building a reproducibility network. PLOS Biology. 2021;19(11):e3001461.

